# Clinical Time-to-Event Prediction Enhanced by Incorporating Compatible Related Outcomes

**DOI:** 10.1101/2022.01.31.478403

**Authors:** Yan Gao, Yan Cui

**Affiliations:** Department of Genetics, Genomics and Informatics, University of Tennessee Health Science Center, Memphis, TN, 38163, USA; Center for Integrative and Translational Genomics, University of Tennessee Health Science Center, Memphis, TN, 38163, USA; Center for Cancer Research, University of Tennessee Health Science Center, Memphis, TN, 38163, USA

## Abstract

Accurate time-to-event (TTE) prediction of clinical outcomes from personal biomedical data is essential for precision medicine. It has become increasingly common that clinical datasets contain information for multiple related patient outcomes from comorbid diseases or multifaceted endpoints of a single disease. Various TTE models have been developed to handle competing risks that are related to mutually exclusive events. However, clinical outcomes are often non-competing and can occur at the same time or sequentially. Here we develop TTE prediction models with the capacity of incorporating data of compatible related clinical outcomes. We test our method on real and synthetic data and find that the incorporation of related auxiliary clinical outcomes can: 1) significantly improve the TTE prediction performance of convention Cox model while maintaining its interpretability; 2) further improve the performance of the state-of-the-art deep learning based models. While the auxiliary outcomes are utilized for model training, the model deployment is not limited by the availability of the auxiliary outcome data because the auxiliary outcome information is not required for the prediction of the primary outcome once the model is trained.

## Introduction

With the rapid advances in health informatic technologies, comprehensive biomedical datasets with multiple related clinical outcomes have become increasingly common. The related clinical outcome variables may represent comorbidity or multifaceted endpoints of a single disease. The multifaceted endpoints provide the measurement of the disease outcomes from different perspectives. The Cancer Genome Atlas^1^ (TCGA) clinical dataset^2^, as a well-known example, includes four variables to characterize the cancer outcome endpoints: overall survival, disease-specific survival, disease-free interval, and progression-free interval. Studies of comorbidity, the coexistence of multiple diseases or disorders in relation to a primary disease in a patient^3^, may also generate multiple related clinical outcome data such as the cardiovascular disease related outcomes in the Sleep Heart Health Study (SHHS) data^4,5^. These related clinical outcomes contain rich information from multiple aspects of disease progression and may stem from common molecular mechanisms or environmental factors.

The Cox proportional hazards (CPH)^6–8^ model has been widely used for time-to-event (TTE) data analysis for decades. In the recent years, deep learning (DL) based methods have been developed for disease classification, diagnosis, and prognosis from personal biomedical data^9–16^. The DL-based TTE models generally outperformed the conventional CPH and other survival analysis models in recent studies^17–23^. However, the black-box nature of deep neural networks makes the DL-based models lack the interpretability that is critical for clinical applications. In these TTE prediction models, a single outcome variable was used as the prediction target, even if multiple related clinical outcome data were available. Developing models to utilize the information from the related outcomes may provide an effective approach to enhance TTE prediction. Various multi-task machine learning models have been developed to handle competing risks that are related to mutually exclusive events^24–28^. However, the widely available compatible related outcomes that may occur at the same time or sequentially have not been formally utilized in the TTE models to improve prediction accuracy.

In this study, we develop a new method, the related outcome incorporator (ROI), to improve the TTE prediction performance for the primary outcome by incorporating related clinical outcome (the auxiliary outcome) data during model training. However, the auxiliary outcome information is not required for the prediction of the primary outcome after model training. Thus, the model application would not be limited by the availability of the auxiliary outcome data. We integrated the ROI with the conventional CPH model and the DL-based model. Using prognosis tasks assembled from real and synthetic datasets, we show that ROI can significantly improve the prediction performance of CPH model while maintaining its interpretability and the ROI can further improve the prediction performance of the state-of-the-art DL-based model.

## Results

### TTE prediction experiments on TCGA data

We compared the TTE prediction models with and without the ROI using the TCGA data. The TCGA dataset contains 4 clinical outcome endpoints for 40 types and categories of cancers (Supplementary Table 1): overall survival (OS), disease-specific survival (DSS), disease-free interval (DFI), and progression-free interval (PFI). We assembled 160 time-to-event prediction tasks using the TCGA clinical data and protein expression data. We filtered out the tasks with less than 50 patients or less than 20 observed events. We further filtered out the tasks with an event-to-censored ratio (E/C) less than 0.2. Finally, we removed the tasks with concordance-index (C-index) less than 0.6 from the baseline CPH model, resulting in 19 tasks from 9 cancer types/categories and 4 different outcomes (**Table 1**). We tested four different TTE prediction models on each task: CPH, CPH with ROI (CPH_ROI), CPH integrated with a deep neural network (CPH_DL), and CPH_DL with ROI (CPH_DL_ROI). The models are described in the Method section. OS and DSS are survival outcomes and are used as a pair of related outcomes for the ROI. Also, PFI and DFI are disease progression outcomes and are used as another pair of related outcomes.

**Table 1.** TTE prediction model performance comparison on TCGA data

| Cancer Type* | Primary Outcome | Auxiliary Outcome | Censored | Events | E/C | C-Index |  |  |  |
| --- | --- | --- | --- | --- | --- | --- | --- | --- | --- |
|  |  |  |  |  |  | CPH | CPH_ROI | CPH_DL | CPH_DL_ROI |
| GBMLGG | OS | DSS | 391 | 274 | 0.70 | 0.74 | **0.79** | 0.80 | **0.81** |
| GBMLGG | DSS | OS | 397 | 246 | 0.62 | 0.74 | **0.80** | **0.81** | **0.81** |
| GBMLGG | PFI | DFI | 325 | 340 | 1.05 | 0.72 | **0.76** | **0.77** | 0.76 |
| KIPAN | OS | DSS | 548 | 206 | 0.38 | 0.71 | **0.73** | 0.73 | **0.74** |
| KIRC | DSS | OS | 364 | 104 | 0.29 | 0.70 | **0.72** | 0.71 | **0.74** |
| KIPAN | DSS | OS | 609 | 133 | 0.22 | 0.70 | **0.76** | 0.78 | **0.80** |
| PanGyn | PFI | DFI | 622 | 459 | 0.74 | 0.69 | **0.70** | **0.70** | 0.69 |
| PanGyn | DFI | PFI | 479 | 216 | 0.45 | 0.67 | **0.70** | 0.71 | **0.73** |
| KIPAN | PFI | DFI | 539 | 213 | 0.40 | 0.67 | **0.70** | **0.72** | **0.72** |
| LGG | OS | DSS | 330 | 98 | 0.30 | 0.66 | **0.73** | 0.75 | **0.76** |
| CESC | PFI | DFI | 141 | 32 | 0.23 | 0.64 | **0.70** | 0.61 | **0.64** |
| LGG | DSS | OS | 333 | 89 | 0.27 | 0.63 | **0.72** | 0.76 | **0.77** |
| PanGyn | OS | DSS | 693 | 388 | 0.56 | 0.63 | **0.64** | 0.66 | **0.67** |
| KIRC | OS | DSS | 312 | 166 | 0.53 | 0.62 | **0.69** | 0.68 | **0.70** |
| PanGyn | DSS | OS | 734 | 313 | 0.43 | 0.62 | **0.67** | 0.68 | **0.70** |
| COADREAD | OS | DSS | 383 | 104 | 0.27 | 0.62 | **0.64** | 0.63 | **0.64** |
| LUAD | DSS | OS | 244 | 80 | 0.33 | **0.61** | 0.52 | 0.53 | **0.58** |
| PanGI | DFI | PFI | 376 | 80 | 0.21 | **0.61** | 0.58 | 0.54 | **0.64** |
| KIRC | PFI | DFI | 322 | 155 | 0.48 | 0.60 | **0.65** | 0.68 | **0.69** |
| Mean |  |  | 428 | 195 | 0.44 | 0.66 | **0.69** | 0.70 | **0.72** |
| Median |  |  | 383 | 166 | 0.40 | 0.66 | **0.70** | 0.71 | **0.72** |

\*See Supplementary Table 1 for the annotations of cancer type abbreviations.

In our experiments, the CPH_ROI outperformed CPH on 17 out of the 19 tasks and CPH_DL_ROI outperformed CPH_DL on 15 out of the 19 tasks and tied with CPH_DL on 2 tasks. The use of ROI improved the mean C-index for the convention Cox model (CPH) by 6.1%, and the mean C-index for the deep Cox model (CPH_DL) by 2.9%. The p-values indicate that the performance improvements by ROI are statistically significant (**Fig.1**).

**Fig 1.**
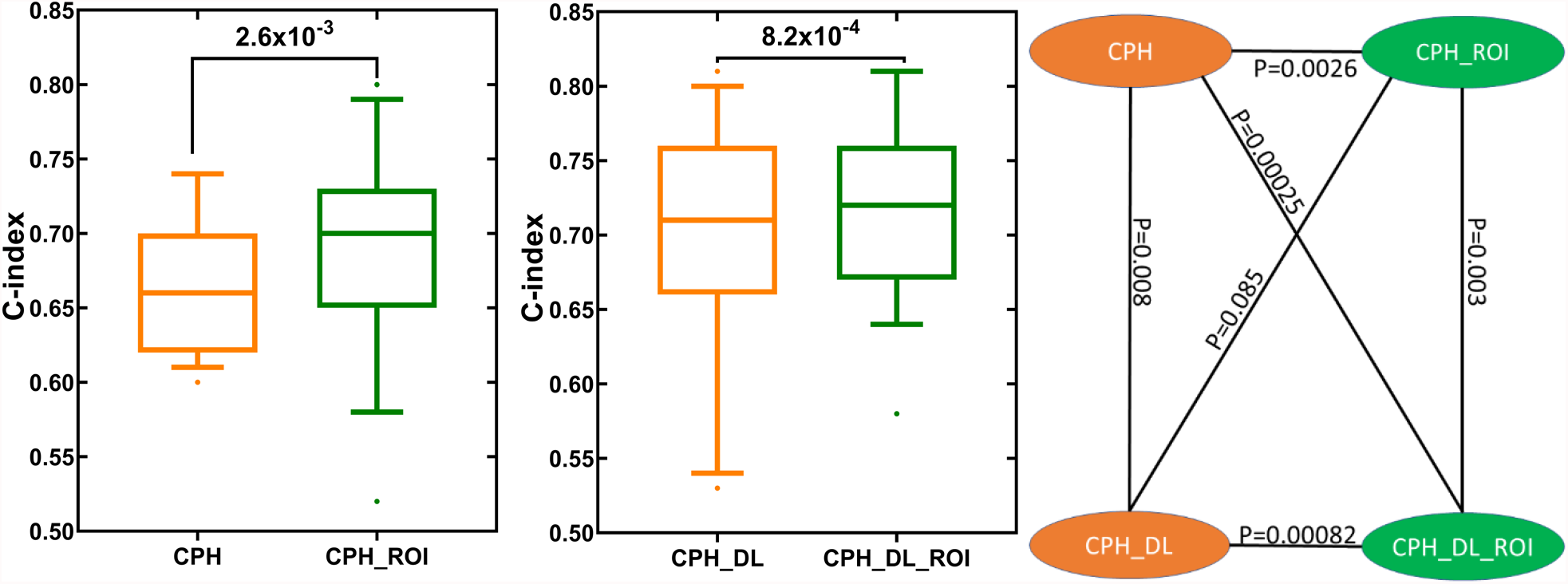
TTE prediction performance comparison between models with and without the related outcome incorporator (ROI). Each box plot shows the distribution of C-index values for the 19 TTE prediction tasks (Table 1). The p values were calculated using one-sided Wilcoxon signed-rank test.

### TTE prediction experiments on SHHS data

To further test the ROI method, we assembled TTE prediction tasks using the data from Sleep Heart Health Study (SHHS). The SHHS dataset contains 5804 samples from 10 different types of cardiovascular disease (CVD), and each sample comprises 1279 clinical features, as well as the follow-up event time. We used Angina as the primary outcome, and Congestive Heart Failure (CHF) and Stroke as the auxiliary tasks are (Table 2). In these experiments, the CPH_ROI model outperformed the CPH model, and the CPH_DL_ROI model outperformed the CPH_DL model. The results show that using ROI can improve the TTE predictions.

**Table 2.** TTE prediction model performance comparison on SHHS data

| Primary outcome | Auxiliary outcome | C-Index |  |  |  |
| --- | --- | --- | --- | --- | --- |
|  |  | CPH | CPH_ROI | CPH_DL | CPH_DL_ROI |
| Angina | CHF | 0.49 | **0.60** | 0.61 | **0.64** |
| Angina | Stroke | 0.49 | **0.59** | 0.59 | **0.62** |

### TTE prediction experiments on synthetic data

To further study our method and test the performance improvement of ROI, we performed an experiment on a synthetic dataset. The synthetic dataset has a total of 300 patients, 200 patients with observed events, and 100 patients with censorship. Each patient has 150 covariates to simulate the high-dimensional biomedical features. The time to the primary outcome (Outcome1) event and auxiliary outcome (Outcome2) event for each patient were generated using the exponential Cox model^29^, in which the hazard function is a linear combination of features *h*(*x*) = *β* × *x*. To simulate the relevance between Outcome1 and Outcome2, we set the weights for their hazard functions have a correlation value greater than 0.8. The results for the primary outcome and the auxiliary outcome are shown in Table 3. We observed a significant improvement in the performance of the models with ROI (CPH_ROI and CPH_DL_ROI) compared to the corresponding models without ROI (CPH and CPH_DL). The results on the synthetic data show that incorporating a related outcome can improve time-to-event predictions, which is consistent with the observations from the real datasets.

**Table 3.** TTE prediction model performance comparison on synthetic data

| Primary Outcome | Auxiliary Outcome | C-Index |  |  |  |
| --- | --- | --- | --- | --- | --- |
|  |  | CPH | CPH_ROI | CPH_DL | CPH_DL_ROI |
| Outcome1 | Outcome2 | 0.67 | **0.81** | 0.79 | **0.84** |
| Outcome2 | Outcome1 | 0.63 | **0.78** | 0.78 | **0.80** |

## Discussion

We developed a method to improve TTE prediction by incorporating related clinical outcomes (ROI) in model training. We tested the utility of the ROI method in two types of models: the conventional CPH model and the deep learning based model that integrates deep neural network and Cox regression (CPH_DL). Consisting with previous observations^17–23^, the DL structures can generate better feature representations and therefore are able to improve the TTE predictions. However, the low interpretability of DL-based models is a major obstacle to their application in healthcare where user (patients and clinicians) trust is critical^30–32^. The use of ROI can improve the TTE prediction performance of the CPH model while maintain its full interpretability. In our experiments on the real and synthetic datasets, the use of ROI improved the prediction performance of the CPH model to the level that is comparable to that of the DL-based model (CPH_ROI vs CPH_DL, Table 1–3). This interesting observation suggests that the use of ROI with CPH model and the integration of DL with CPH model may lead to a comparable level of prediction performance improvement. The use of ROI also further improved the performance of the DL-based model. Time-to-event analysis is widely used in many areas beyond medicine, including engineering, economics and finance. We expect that the ROI framework may also be applicable to TTE predictions in more broader areas.

## Methods

### Datasets

We used the TCGA clinical and protein expression datasets from the Genome Data Commons (GDC, https://gdc.cancer.gov). We filtered the proteins with missing values, resulting in 189 protein features. Patients with missing values for follow-up time or event indicators were removed. For each survival analysis task, the protein expression values were standardized by removing the mean and scaled to unit variance. The clinical outcome is the time in days and the event indicator for endpoints: OS, PFI, DFI, and DSS.

For the Sleep Heart Health Study (SHHS)^4,5^ dataset, we acquired the data from the National Sleep Research Resource (https://sleepdata.org/datasets/shhs). We removed the records with 20% missing features and dropped the features with missing values. We converted the *rcrdtime* (total recording time) into the number of minutes and filtered out samples without follow-up time information. After the preprocessing, we got 1514 uncensored events from 10 outcomes, and each record has 374 features. For each event, we applied an unsupervised feature selection method to select 150 features with top mean absolute deviation values. We then filtered out the clinical outcomes with less than 50 samples. The final dataset contains three clinical outcomes: Angina (121 patients), Congestive Heart Failure (197 patients), and Stroke (79 patients).

### The baseline model

In our study, for each survival analysis task, the baseline model we used is the Cox proportional-hazards (CPH) model^33^ (Fig 2A) implemented in the python lifeline package^34^. To improve the robustness of the model, we set the penalty weight of 0.0001 to the coefficients during fitting. All other parameters remain unchanged.

**Fig 2.**
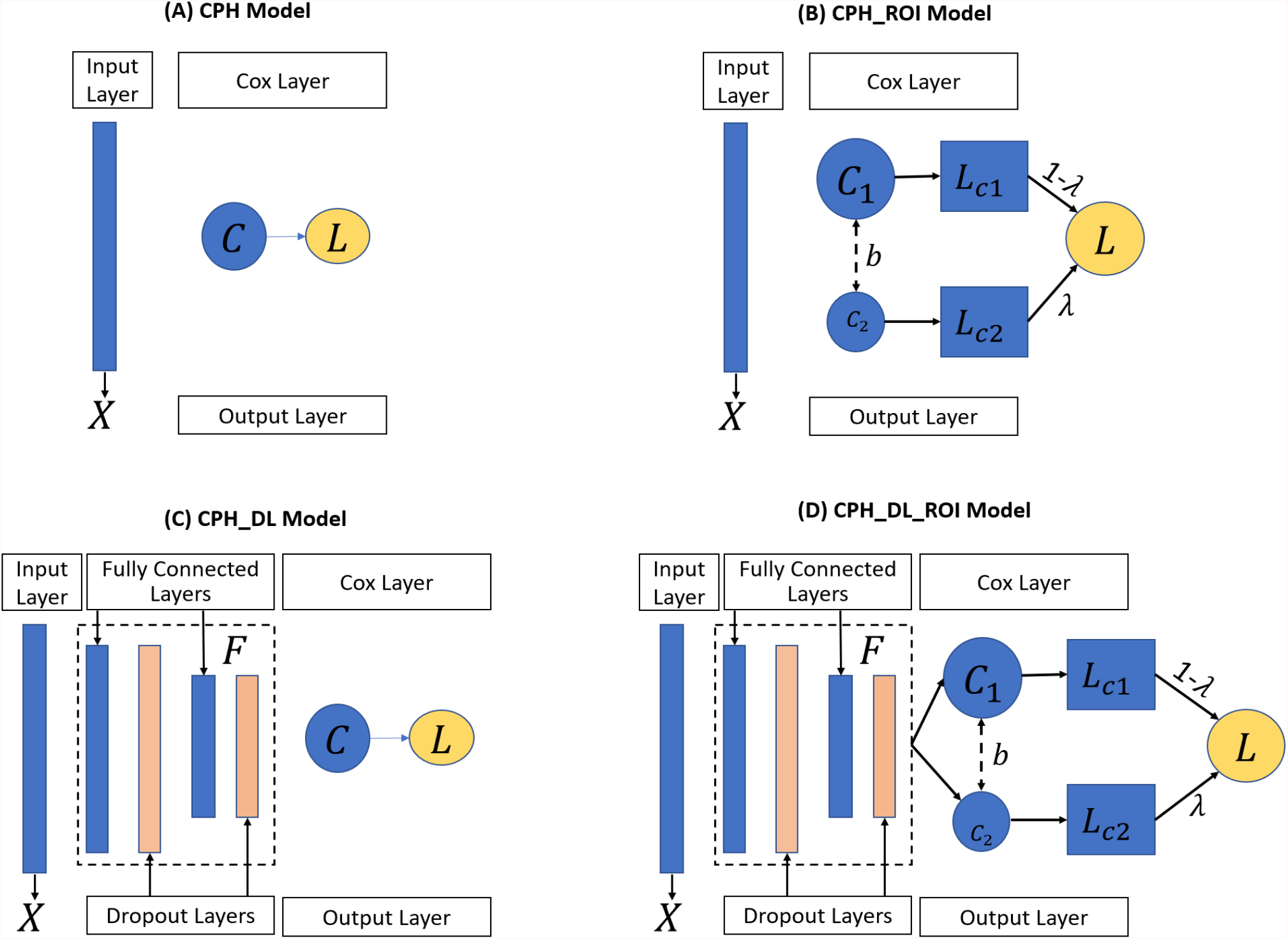
The four TTE prediction models: (A) CPH Model, (B) CPH_ROI Model, (C) CPH_DL Model and (D) CPH_DL_ROI Model. *X* represents the input features, *C* is the Cox regression layer, *L* is the partial hazard loss function, *λ* is the weight to balance outcomes, and *F* is the feature extractor to map the input feature into an embedding space. **CPH**: Cox proportional hazards. **ROI**: related outcome incorporator. **DL**: deep learning.

### The CPH_ROI model

The proposed CPH_ROI model has two parts, and each part is a regularized CPH model (Fig. 2B). The loss function of CPH_ROI (Equation 1) is a linear combination of two related events, the loss of the main task and the loss for the related task. In the equation, *λ* is a hyperparameter used to balance the loss between the primary outcome and the auxiliary outcome, *β*_1_ and *β*_2_ are the log partial hazard ratio for the main and the auxiliary outcomes, *b* is the bias, *U*_1_ and *U*_2_ are the set of uncensored patients for the main and the auxiliary outcomes. In equation 2, 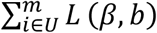 is the log partial likelihood which is defined in Equation 3, *U* is the set of uncensored patients, and *λ*_1_ is the weight of the *L*_2_ regularization term. In Equation 3, *m* is the number of patients, *R*(*T*_*i*_) is the risk group in which each patient’s survival time is greater than *T*_*i*_, *δ*_*i*_ is the event status of patient *i, δ*_*i*_ = 1 if an event (like death) occurred, and *δ*_*i*_ = 0 when there is a censorship. The *h*(*X*_*i*_, *β, b*) in equation 4 is the linear hazard function, where *X*_*ij*_ is the *j*^*th*^ feature of patient *i, β*_*i*_ is *j*^*th*^ weight of *β*, and *b* is the bias. During the training, we set *λ* to 0.2, the parameters of *β*_1_, *β*_2_, *b* were randomly initialized and optimized by minimizing the objective function in equation 1. Adopting the idea of transfer learning for high-dimensional linear regression^35^, we set the two linear functions to share the same bias.

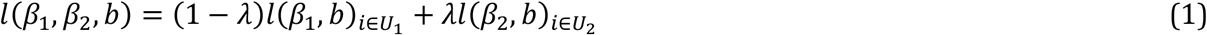

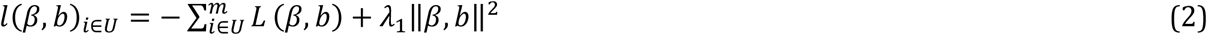

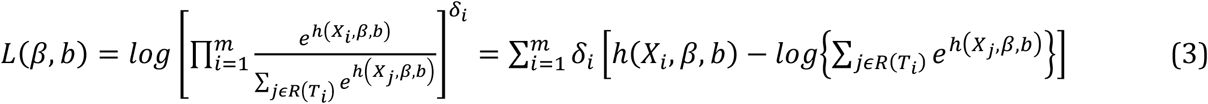

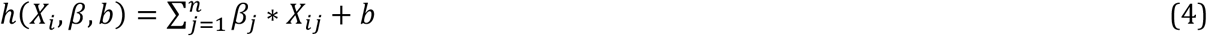

### The CPH_DL model

In the CPH_DL model, a Cox regression layer was added on top of a deep neural network structure (Fig. 2C). The loss function is shown in Equation 5, which has two parts: a deep neural network-based feature extractor *F* and a Cox regression layer *C*, where *F* maps the high-dimensional input feature into an embedded space *Z*, and then *C* makes a prediction from *Z*. In Equation5, *λ*_1_ is the *L*_2_ regularization weight, *W* is the weight of *F, β* and *b* are the log partial hazard ratio and the bias of *C*. All other notions are the same as the CPH_ROI model. The feature extractor *F* is a 4-layer deep neural network structure: a fully connected (FC) layer of 100 nodes followed by a dropout layer (with p=0.5), an FC layer of 50 nodes followed by a dropout layer (with p=0.5). These two dropout layers and the regularization term are used to avoid overfitting during training.

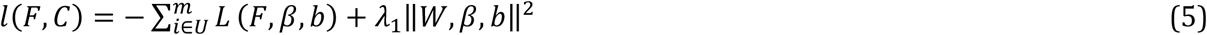

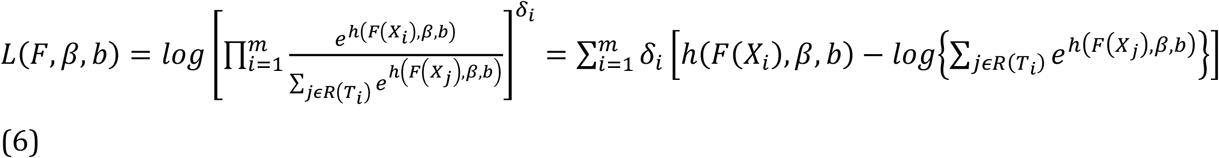

### The CPH_DL_ROI model

The CPH_DL_ROI model (Fig. 2D) incorporates the related outcome in its loss function (Equation 7), where *λ* is the adjustment weight, *F* is the feature extractor, *C*_1_ and *C*_2_ are the Cox regression layer for each outcome. Similar to the CPH_ROI model, we set *C*_1_ and *C*_2_ share the bias and set *λ* to 0.2. The notion of *l*(*F, C*), *U*_1_ and *U*_2_ are same as the notions defined in the CPH_ROI and CPH_DL models.

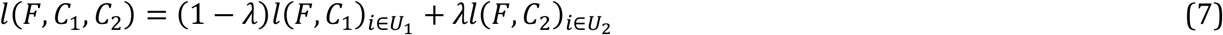

### Experiment setup

For a given task with two related outcomes 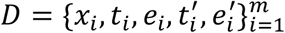, *r* is the number of patients, and for patient *i, x*_*i*_ is the input feature, *t*_*i*_ and *e*_*i*_ are the event time and status of the primary outcome,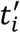 and 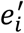 are the time and event for the auxiliary outcome. We formulate it as *D* = {*D*_1_ ∪ *D*_2_}, where 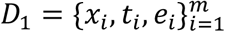 is the main task and 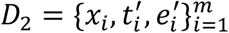 is the auxiliary task. For each task *D*_*i*_, we applied a stratified ten-fold cross-validation for training and testing splitting. The dataset was stratified by the event status to ensure a uniform distribution of events across each fold. In each fold, the training set of *D*_1_ was used to train a CPH and CPH_DL models, while the training sets of *D*_1_ and *D*_2_ were used to train the CPH_ROI and CPH_DL_ROI models. For the CPH_ROI and CPH_DL_ROI models, in the testing stage, only the input features from the testing set of *D*_1_ were used to calculate the risk of each sample, and these risk scores were used for evaluation.

### Evaluation metric

We evaluated the prediction performance of each model using the concordance index (C-index)^36^, which measures the proportion of concordant pairs among the total number of possible pairs. Those pairs were discarded if the earlier time is censored. For a testing set, its risk scores with event time and event status were used to calculate the C-index. A higher C-index value indicates a better TTE prediction model. A C-index of 1.0 indicates a perfect prediction model, while a C-index of 0.5 indicates a totally random prediction model.

### Synthetic data generator

We developed a synthetic data generator to simulate the dataset for our simulation study. The simulated dataset can be formulated as 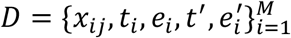, where *M* is the total number of patients, *x*_*ij*_ is the *j*^*th*^ input feature of patient *i, t*_*i*_ and *e*_*i*_ are the event time and status of the primary outcome,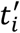 and 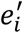 are the event time and statues of the auxiliary outcome of patient *i*. We simulated the feature matrix *x*_*ij*_ from a uniform distribution with the lower boundary −1 and higher boundary of 1. For each patient, we set the feature size to 150 to simulate the high-dimensional biomedical data. The main event time for patient *i* was simulated using the exponential Cox model, as shown in Equation 8, where *λ* is the baseline function, *EXP* is an exponential distribution with a *mean* = 3000, and *β*_*i*_ is the weight of feature *j* which was generated from a uniform distribution. With the simulated event time, we set a cut-off time threshold to simulate the “end-of-study” to keep a portion of desire uncensored samples in the dataset.

To simulate the event time and status of the related outcome, we added a random noise to the weight of the primary outcome, 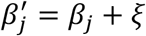, where *ξ* is a uniform distribution with a range of 0 and 1. We control the correlation between *β* and *β*^*′*^ to be greater than 0.8 to ensure the relevance of two outcomes. For both events, we generated a total of 300 patients, 200 patients with observed events, and 100 patients with censorship.

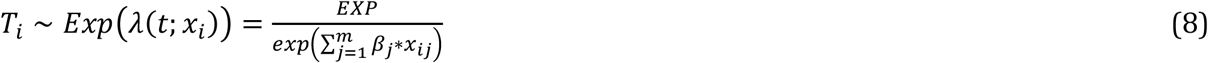

## Data availability

The TCGA data was downloaded from the Genome Data Commons (https://gdc.cancer.gov/about-data/publications/pancanatlas). The SHHS data was downloaded from the National Sleep Research Resource website (https://sleepdata.org/datasets/shhs).

## Acknowledgements

The machine learning experiment tasks in this work were assembled from TCGA data generated by TCGA Research Network (https://www.cancer.gov/tcga), and The SHHS data generated by Sleep Heart Health Study (https://sleepdata.org/datasets/shhs). This work was supported by the NIH grant R01CA262296 and The UT Center for Integrative and Translational Genomics.

## Notes

### Competing Interest Statement

The authors have declared no competing interest.

